# Predominant Patterns of Splicing Evolution on Human, Chimpanzee, and Macaque Evolutionary Lineages

**DOI:** 10.1101/204255

**Authors:** Jieyi Xiong, Xi Jiang, Angeliki Ditsiou, Yang Gao, Jing Sun, Elijah D. Lowenstein, Shuyun Huang, Philipp Khaitovich

## Abstract

Although splicing is widespread and evolves rapidly among species, the mechanisms driving this evolution, as well as its functional implications, are not yet fully understood. We analyzed the evolution of splicing patterns based on transcriptome data from five tissues of humans, chimpanzees, rhesus macaques, and mice. In total, 1,526 exons and exon sets from 1,236 genes showed significant splicing differences among primates. More than 60% of these differences represent constitutive-to-alternative exon transitions while an additional 25% represent changes in exon inclusion frequency. These two dominant evolutionary patterns have contrasting conservation, regulation, and functional features. The sum of these features indicates that, despite their prevalence, constitutive-to-alternative exon transitions do not substantially contribute to long-term functional transcriptome changes. Conversely, changes in exon inclusion frequency appear to be functionally relevant, especially for changes taking place in the brain on the human evolutionary lineage.

## INTRODUCTION

Alternative splicing (AS) enables the expansion of the transcriptome repertoire through the creation of novel transcript isoforms (1). The proportion of genes undergoing AS increases with organismal complexity, from 25% in nematode worms to 95% in humans (2,3). Alternative splicing further increases transcriptome variability by creating isoform variation among tissues within an individual (3,4), among different age groups (5), and among human populations (6). Comprehensive comparison studies in vertebrates (7) and mammals (8) revealed that splicing divergence increases rapidly as phylogenetic distances increase.

Numerous studies reported that AS plays a critical role in various physiological processes in eukaryotes (9–11). Yet, functions of the vast majority of alternative isoforms are still unknown. A large proportion of detected isoforms might not be functional, and may instead represent aberrant splicing products, as indicated by their low expression and low splice site sequence conservation (12). Similarly, evolutionary analysis of splicing patterns among primate species indicates that the majority of observed splicing differences might represent evolutionary neutral drift (13).

Previous studies also suggest that evolutionary dynamics, mechanisms, and functionality of alternatively spliced isoforms may differ substantially depending on their splicing patterns, inclusion frequency, and evolutionary history. For example, novel isoforms formed by exonization may not be as functional as those produced by other types of exon recruitment, independent of novel exon inclusion frequency (14).

In this study, we investigated the relative frequency, evolutionary conservation, evolutionary birth and death rates, as well as potential functionality of isoforms created by different mechanisms. Our analysis was based on transcriptome sequencing data collected from five somatic tissues from humans, chimpanzees, rhesus macaques, and mice. The relatively short divergent times (15–18) and large proportion of alternatively spliced genes in these species provide a good system to investigate the different types and properties of splicing changes along the evolutionary lineages.

## RESULTS

### Splicing differences among species

We estimated splicing differences among humans, chimpanzees, rhesus macaques, and mice using poly-A-plus transcriptome data collected in three brain regions (prefrontal cortex, visual cortex, and cerebellum), kidney, and muscle in six adult individuals of each species (Supplementary Table S1) (19). The total transcriptome data contained 1.87 billion 100 nucleotide (nt) long reads, 1.34 billion (72%) of which were mapped to the corresponding genome (Supplementary Table S2).

In the first step of our splicing analysis we constructed consensus transcriptome maps of the three primate species (human, chimpanzee, and macaque) or all four species (primates and mouse), based on the RNA-seq data. Specifically, after mapping RNA-seq reads to the corresponding genome in an annotation-free manner, we mapped all identified canonical splice junctions to the human genome. This procedure yielded 224,325 splice junctions detected in three primate species and 94,544 splice junctions in all four species. Using these junctions we identified exon sets consisting of an internal gene segment and two flanking exons. Exon sets containing low occurrence isoforms (inclusion < 10% in all individuals) were excluded from further analysis. Following this procedure we detected 5,480 cassette exon sets with sufficient exon junction coverage in three primate species and 2,859 cassette exon sets in all four species.

Splicing variation analysis, conducted based on percent-spliced-in (PSI) measurements of internal gene segments (single exons or exon groups) located within the 1,925 exon sets detected in all tissues of three primate species, showed clear separation among organs and species (Figure 1A). Concordantly, 23% of the total splicing variation was explained by the differences among tissues and 18% by the differences among the three primate species (permutations, p<0.0001) (Figure 1B). Similar results were observed using the four species data (Figure 1C, D; Supplementary Figure S1).

**Figure 1.**
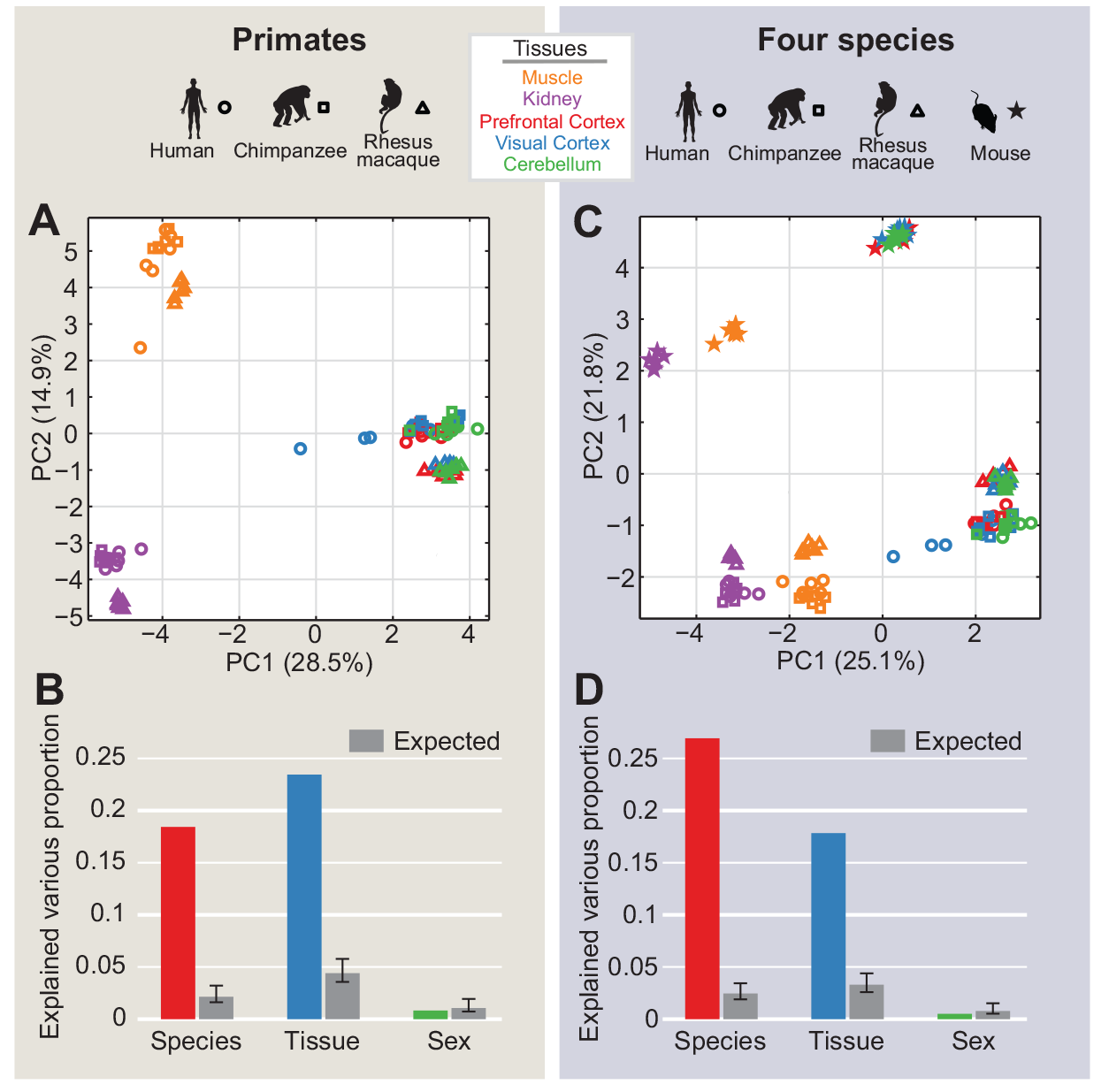
Overview of splicing variation. (A, C) PCA plots based on PSI values for 5,480 events in 90 primate samples (A) and 2,859 events in 120 samples from the four species (C). The symbols show species, and colors show tissues. (B, D) The proportion of total splicing variation explained by the factors species, tissues and sex in primates (B) and four species (D). The proportion of variation explained by chance (gray bars) and their 95% confidential intervals (error bars) were calculated by 1,000 permutations of the factor labels.

To assess the validity of identified splicing differences we compared them to published microarray and PCR results (20). Of the 281 and 340 splicing differences reported in cerebellum between humans and chimpanzees or macaques respectively, 111 and 127 were covered by exon sets detected in our study. Of these, 94 (84.7%) and 103 (81.1%) showed consistent difference direction in our data (Supplementary Figure S2). These proportions are close to the RT-PCR validation rate (83%) reported in the study (20). Direct comparison between our data and the published PCR validation tests showed that 20 of the 23 detected splicing events showed consistent differences among three primates, while the remaining three did not yield conclusive results based on visual inspection of the gel images (Supplementary Table S3).

### Patterns of splicing evolution

To identify significant splicing differences among species we applied the generalised linear model to 5,480 cassette exon sets detected in primates and to 2,859 cassette exon sets detected in four species. This was done assuming that the distribution of junction-spanning reads followed the quasi-binomial model (21–23). The false discovery rate was set to 5% based on 1,000 permutations of species labels. We further required that the mean PSI difference between species be greater or equal to 0.2. Based on these criteria, a total of 1,526 cassette exon sets from 1,236 genes showed significant splicing differences among primates in at least one tissue. Similarly, 1,385 cassette exon sets from 1,035 genes showed significant splicing differences among the four species in at least one tissue.

In agreement with previous reports (7,8), the number of splicing differences between species was mainly proportional to the genetic distances (Pearson correlation, r=0.88, p<0.0001; Figure 2A, Supplementary Table S4).

**Figure 2.**
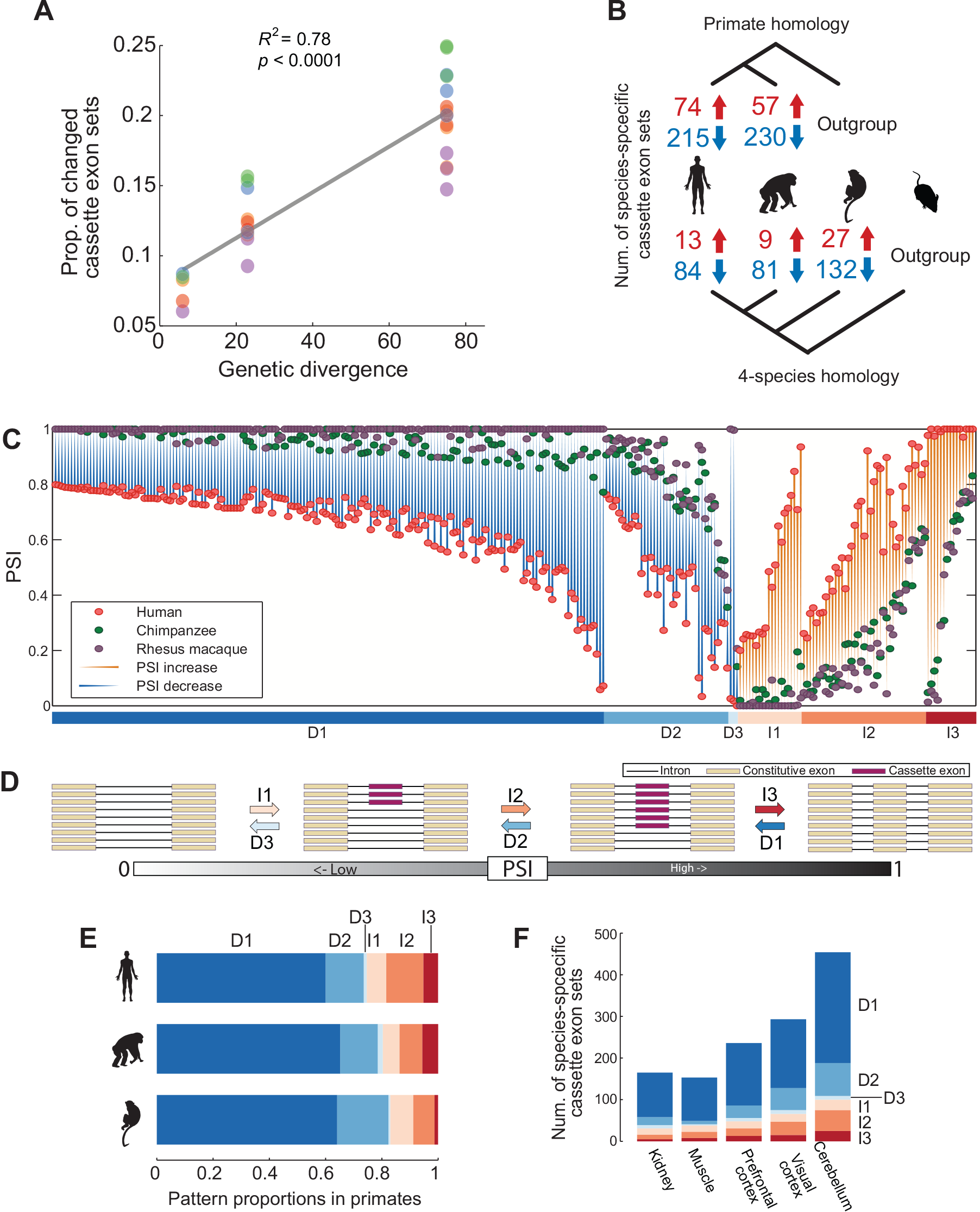
Splicing divergence among species. (A) The relationship between splicing divergence and genetic distances among species. The colors represent tissues, as in Fig 1. (B) The number of cassette exon sets showing splicing changes specific to the lineages detected among primate species (top) or in the four species comparison (bottom). The red and blue numbers represent the number of cassette exon sets showing increased and decreased exon inclusion frequencies respectively. (C) Human-specific splicing changes. When splicing changes were detected in multiple tissues, the one with largest PSI difference is shown. The colored bar marks the six evolutionary splicing patterns. The same color-coding of splice patterns is used for the arrows in (D), and bar plots in (E) and (F). (D) Schematic representation of the six evolutionary splicing patterns. (E) The proportions occupied by each of the six splice patterns on the three primate lineages. (F) The number of cassette exon sets with species-specific splicing in each of the six patterns in each of the five tissues.

Assignment of splicing differences to the evolutionary lineages and identification of the direction of splicing changes produced an unexpected observation: on all three evolutionary lineages changes leading to less exon inclusion were on average more than four times more frequent than changes leading to greater exon inclusion (Figure 2B). To understand the possible evolutionary mechanisms of this phenomenon, we further sorted splicing events into six evolutionary patterns. The three patterns of decreased exon inclusion are: (D1) decreased inclusion of previously constitutive exons (constitutive-to-alternative), (D2) decreased inclusion of alternative exons (alternative-to-alternative), and (D3) constitutive exclusion of alternative exons (exon death). The three patterns of increased exon inclusion are: (I1) birth of alternative exons from intronic sequences (exon birth), (I2) increased inclusion of alternative exons (alternative-to-alternative), and (I3) constitutive inclusion of previously alternative exons (alternative-to-constitutive) (Figure 2C-D).

Strikingly, distribution of observed splicing differences among these patterns was far from even. The constitutive-to-alternative D1 pattern contained 78% of all changes leading to a decrease in exon inclusion and 61% of all species-specific splicing events identified on the three primate lineages (Figure 2E). Modifications of the inclusion frequency of alternative exons (alternative-to-alternative D2 and I2 patterns) occupied 15% and 10% of all splicing events, respectively. Exon birth and alternative-to-constitutive patterns I1 and I3 were even more rare, constituting 7% and 5% of all splicing events respectively. The exon death pattern D3 was the most rare, contributing less than 2% of all splicing events. This uneven distribution of splicing patterns was observed in each of the five tissues (Figure 2F, Supplementary Figure S3).

### Evolutionary properties of the main splicing patterns

If changes included in the D1 pattern, *i.e.* constitutive-to-alternative exon transitions, are retained during species evolution, then they should lead to a rapid increase in the proportion of alternative isoforms. To assess this, we examined the evolutionary conservation of splicing changes in the three dominant splicing patterns: D1, D2, and I2. Using mice as an outgroup, we assigned splicing events to the human, chimpanzee, and macaque evolutionary lineages, as well as to the ape lineage leading to the most recent common ancestor of humans and chimpanzees (Figure 3A, Methods).

**Figure 3.**
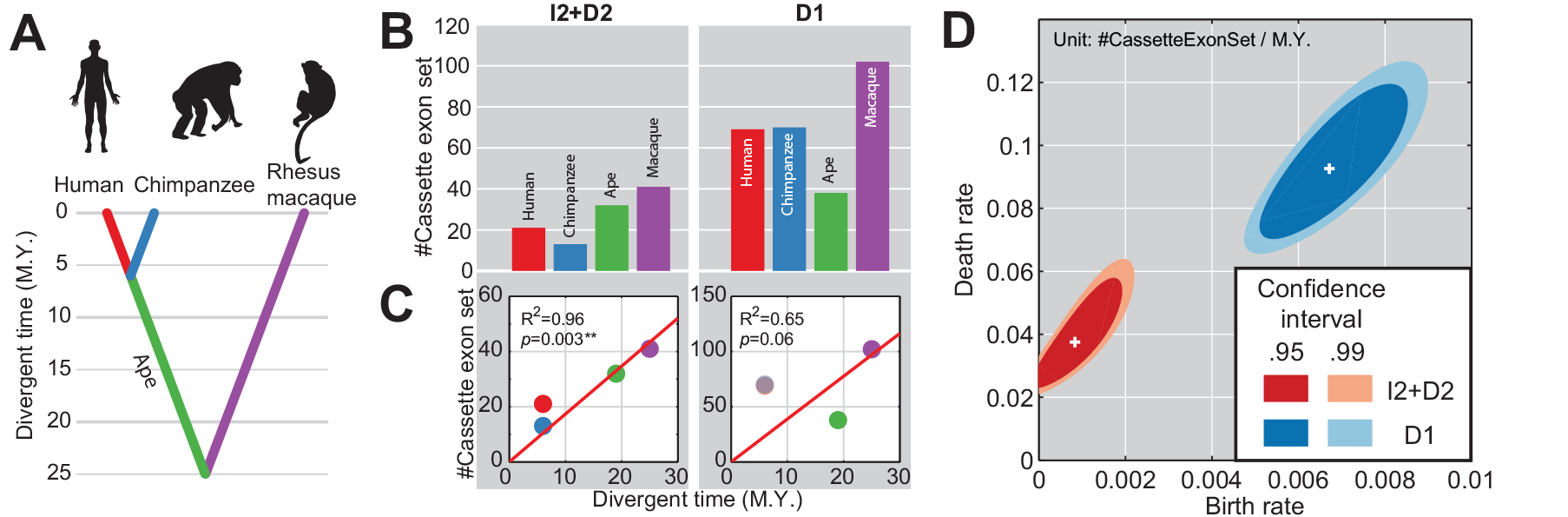
Evolutionary properties of the main splicing patterns. (A) Schematic diagram of the evolutionary tree of the three primate species. The same lineage color-coding is used in (B) and (C). M.Y., millions of years. (B) The number of gene segments within the I2 and D2 patterns (I2 + D2) or within the D1 pattern in four-species. (C) The relationship between species divergence and the number of cassette exon sets for the I2 + D2 patterns and the D1 pattern. In the D1 correlation plot, the dots representing human and chimpanzee values largely overlap. (D) Estimated birth rate (x-axis) and death rate (y-axis) of splicing events in the I2 + D2 patterns (red) and the D1 pattern (blue). White crosses show the maximum likelihood estimates, the dark and light color areas show the 95% and 99% confidence intervals respectively.

Notably, the proportion of D1 splicing events on the ape evolutionary lineage was approximately two times smaller than on either the human or chimpanzee lineages (Figure 3B). This finding is unexpected, as the estimated length of the ape lineage is three to four times greater that that of the human or chimpanzee lineages (15–18). The scarcity of D1 events assigned to the ape lineage can be explained either by the recent independently occurring acceleration of D1 pattern events in the human and chimpanzee lineages or, more likely, by the rapid evolutionary loss of D1 events originating on the ape lineage. In contrast to D1, other splicing patterns such as D2 and I2 have similar evolutionary pattern characterized by more events assigned to longer evolutionary lineages (Figure 3C). This difference suggests that constitutive-to-alternative exon transitions (D1 pattern) appear and disappear more often during the course of evolution than other splicing events.

We next calculated evolutionary birth and death rates of splicing events in the three dominant splicing patterns, D1, D2, and I2, on the human, chimpanzee, ape, and macaque evolutionary lineages using the maximum likelihood approach. We show that in agreement with our predictions, the birth rate of D1 splicing events was eight times greater than the birth rate of D2 and I2 splicing events taken together: 0.0067 (0.0051 - 0. 0085) splicing events per cassette exon set per million years for the D1 pattern and 0. 00083 (3.0×10−6 - 0.0019) for the D2 + I2 patterns. The death rate of these novel splicing events, *i.e.* reverse splicing changes, was also substantially higher for the D1 pattern than for the D2 and I2 patterns: 0.093 (0.071 - 0.12) per event per million years for the D1 pattern and 0.037 (0.023 - 0.058) for the D2 + I2 patterns (Figure 3D). These results further confirm the conclusion that evolutionary events leading to alternative splicing of constitutive exons are common, but the majority of the resulting alternative isoforms are rapidly lost during the course of evolution. By contrast, changes affecting the inclusion frequency of existing alternative exons are less frequent but more likely to be conserved. Thus, even though decreased inclusion of constitutive exons is the most common pattern in splicing evolution, it might not play the leading role in the increase of isoform repertoire over substantial evolutionary time.

### Sequence conservation in the main splicing patterns

To assess the functional significance of splicing events in the three dominant patterns, D1, D2, and I2, we examined the sequence and functional conservation of corresponding alternative exons. For all three patterns, the vast majority of splicing events assessed in our study (86-97%) resided within the protein-coding region (CDS) of a gene (Supplementary Figure S4). Among them, the proportion of splicing events disrupting the reading frame was significantly higher in the D1 pattern (Figure 4A). Furthermore, DNA sequence conservation of alternatively spliced exons, as well as conservation of the entire CDS region, was significantly lower in the D1 pattern (Figure 4B). These observations suggest the neutral or deleterious character of D1 pattern events.

**Figure 4.**
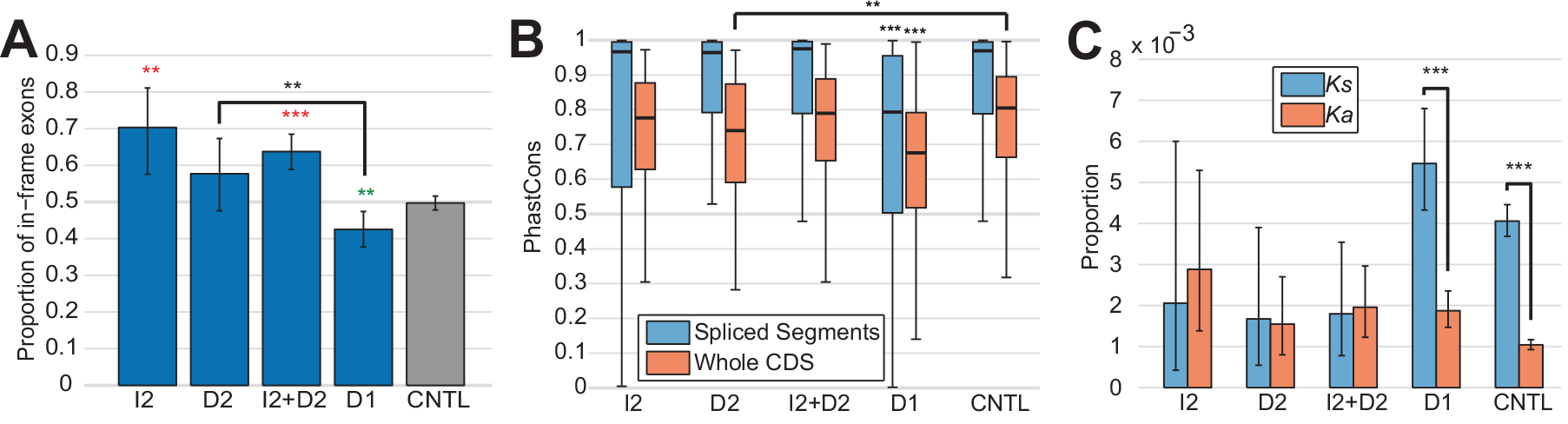
Sequence conservation properties of the main splicing patterns. (A) The proportion of exons preserving the open reading frame in the dominant evolutionary splicing patterns. Red and green asterisks respectively show the significance of higher and lower ratios calculated in comparison with control exons (CNTL) using the two-tailed Fisher's exact test. (B) The distribution of phastCons scores in the dominant evolutionary splicing patterns, for gene segments of differently spliced regions (blue) and for whole coding regions (orange). In both assays, the D1 pattern shows significantly lower sequence conservation for both spliced exons (p<0.0001) and coding regions (p<0.001) compared to any other pattern (two-tailed Wilcoxon test). (C) The rates of synonymous (Ks) and nonsynonymous (Ka) substitutions on the human and chimpanzee evolutionary lineages. In (A, C), the error bars show the 95% confidence intervals.

Surprisingly, despite the relatively high sequence conservation of D2 and I2 exons, they displayed higher Ka/Ks ratios when compared to D1 or control exon groups (Figure 4C). This finding parallels observations of elevated Ka/Ks ratios in alternatively spliced exons of mammals, suggesting a stronger directional selection pressure on these regions (24,25).

### Regulatory mechanisms of the main splicing patterns

The different evolutionary properties of the D1 and D2/I2 patterns suggest differences in their regulatory mechanisms. To assess this, we examined the extent of DNA sequence changes affecting splice site strength, as well as the presence of regulatory motifs in the vicinity of splice sites.

D1 pattern events indeed contained more sequence changes within the 5′-splice sites flanking the alternatively spliced segment compared to control exons (Fisher exact test, p<0.05) (Figure 5A). Furthermore, the resulting changes in splice site strength coincided with changes in exon inclusion frequency (binomial test, p<0.001). This connection between splice site mutation and exon inclusion frequency changes is consistent with the results of a previous comparison between humans and mice (26). However, 5′-splice sites flanking alternatively spliced segments in the D2 and I2 patterns show fewer sequence changes compared to control exons (Fisher exact test, p<0.0005) (Figure 5A). Still, for the few observed sequence changes the resulting decrease or increase in splice site strength also coincided with the change in exon inclusion frequency (binomial test, p<0.05). These results indicate that mutations affecting 5′-splice site strength contribute substantially to splicing changes in all three of the dominant patterns.

**Figure 5.**
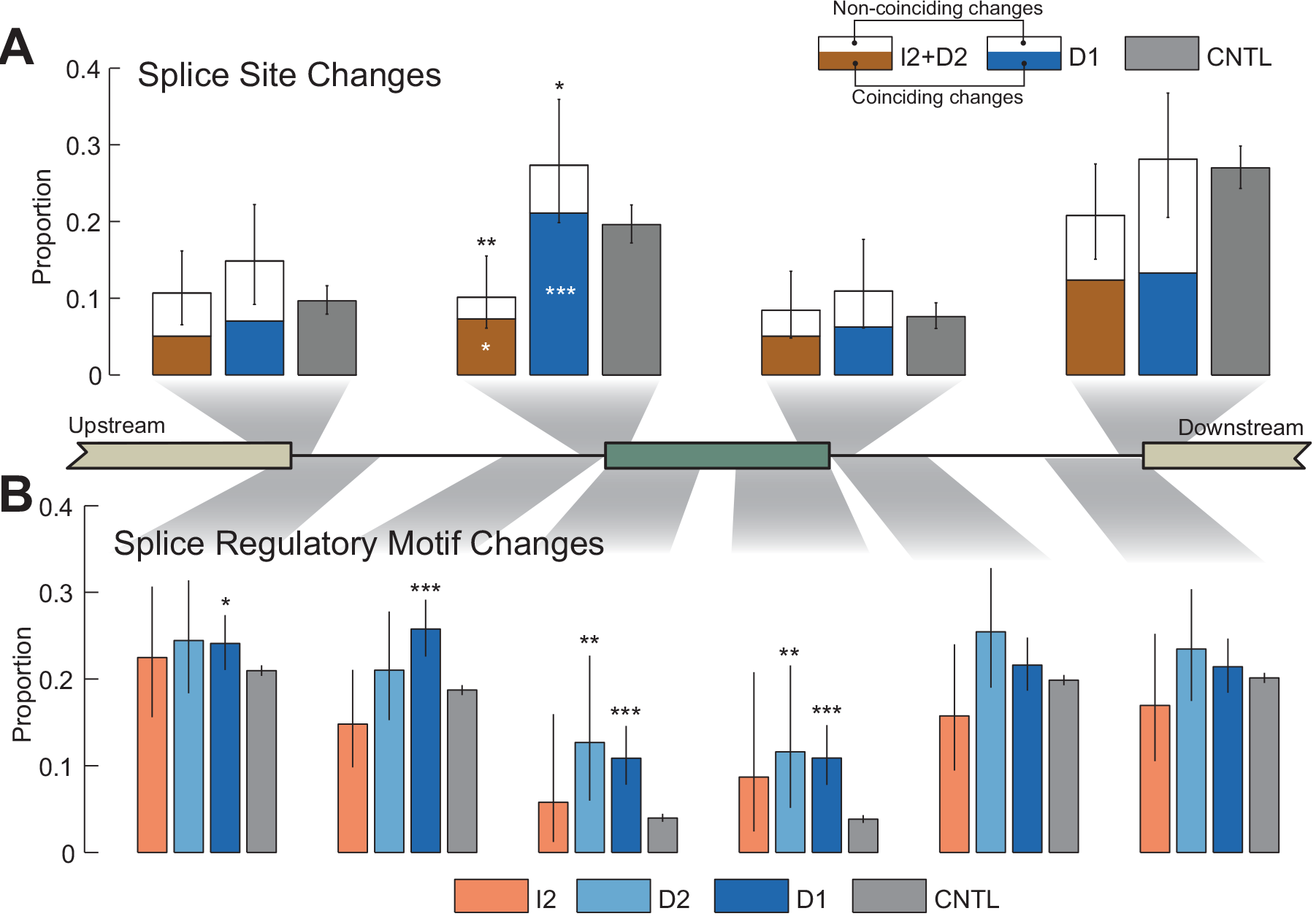
Regulatory properties of the main splicing patterns. (A) The proportion of splice sites with strength changes on the ape and rhesus macaque lineages. The colored and white sections of the bars show the strength of change consistent and inconsistent with the direction of exon inclusion frequency changes, respectively. White asterisks within the colored sections of the bars indicate the significance of the colored proportions calculated using the binomial test. (B) The proportion of regulatory motif turnover on the human and chimpanzee evolutionary lineages. In (A) and (B), error bars show 95% confidence intervals. The schematic exonintron structure in the middle shows the positions of splice sites in (A) and evaluated regions in (B). The length of evaluated regions in introns and exons depicted in (B) were 200nt and 39nt, respectively.

The birth and death frequency of regulatory splicing motifs near junctions flanking alternatively spliced segments also varied among the three dominant splicing patterns. Both D1 and D2 patterns had a higher proportion of sequence changes leading to regulatory motif turnover in the 5′-end region of alternative exons compared to control exons (Fisher exact test, p<0.05) (Figure 5B). For the D1 pattern, further changes affecting regulatory motif turnover occurred in the proximal and distant positions of 5′-introns, a region which has been reported as important for alternative splicing regulation (27) (Fisher exact test, p<0.05). Lastly, no increase in regulatory pattern turnover was detected for the I2 pattern (Figure 5B). These observations indicate that changes in regulatory elements contribute differently to the evolution of the three dominant splicing patterns.

### Human-specific features of splicing evolution

In order to characterize features of splicing evolution specific to humans, we compared the frequencies of lineage-specific splicing events between the human and chimpanzee lineages. In agreement with the phylogenetic distances, splicing changes corresponding to D1 and D2 patterns were distributed equally on the human and chimpanzee lineages in all five tissues. Conversely, splicing changes corresponding to the I2 pattern occurred approximately two times more frequently on the human lineage than on the chimpanzee lineage (one-sided Fisher exact test, p=0.03). This difference was even greater for the exon sets present in all four species (Figure 6A).

**Figure 6.**
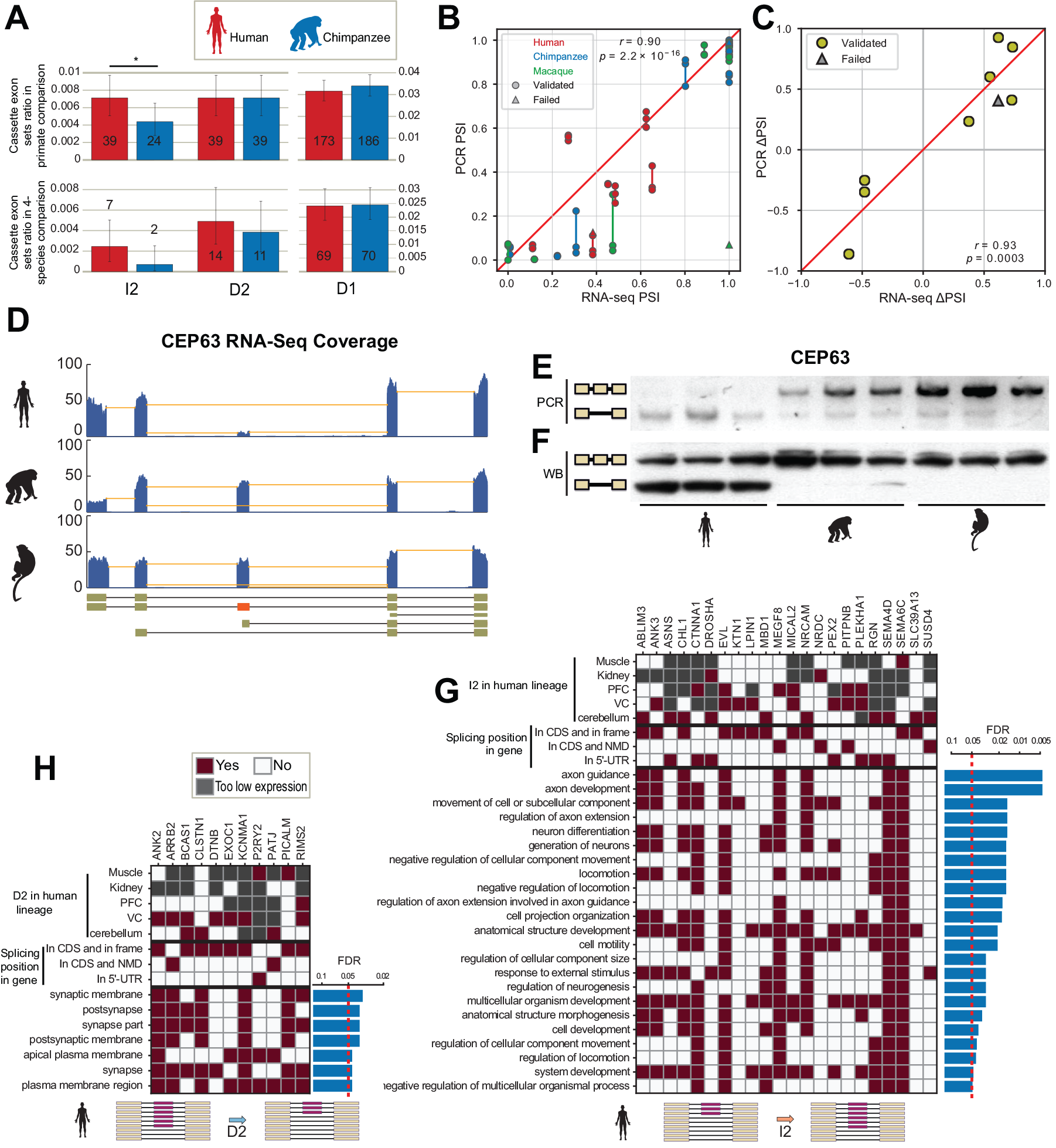
Human-specific features of splicing evolution. (A) The proportion of cassette exon sets showing species-specific splicing changes for each of the dominant splicing patterns. The proportions for human and chimpanzee lineages were calculated in the primate species comparison (upper panel) and by four species comparison (lower panel). Exon set numbers are shown within the bars. The error bars show 95% confidential intervals based on the binomial distribution. The asterisks mark the significance of the ratio differences between the human and chimpanzee branches (one-tailed Fisher's exact test, p<0.05). (B) Comparison of PSI values obtainedbased on RNA-seq and PCR measurements. (C) The comparison of human-specific PSI changes (human minus the mean of chimpanzee and rhesus) was calculated based on RNA-seq and PCR measurements. In (B, C), one case showing inconsistent splicing patterns in macaques in PCR experiment compared to RNA-seq data is shown as triangles. (D-F) RNA-seq reads coverage (D), PCR results (E), and Western blot results (F) for human-specific splicing in the CEP63 gene. In (D), coverage in three primates using visual cortex data is shown as blue blocks; junction reads are shown as orange lines. The y-axis indicates the reads number. Human gene annotation is shown at the bottom of the figure, the differentially spliced exon (located at 930-1067nt within the coding region) is colored in red. In (E), PCR was conducted in visual cortex samples of three individuals for each of the three primate species. In (F), Western blot was conducted in cerebellum samples from three individuals for each of the three primate species. In both (E) and (F), the longer and shorter products corresponding to exon inclusion and exclusion isoforms are detected in the upper and lower rows, respectively. (G, H) Characteristics of genes showing I2-type (G) and D2-type (H) splicing changes on the human evolutionary lineage and the corresponding enriched GO terms. The red color indicates the presence of the feature. Bar plots show the significance of each GO term after multiple test correction.

To further examine the authenticity of human-specific splicing changes, we conducted additional semi-quantitative PCR experiments. We designed primer pairs for twelve out of 289 human-specific splicing events detected using RNA-seq data. Nine of these primer pairs effectively amplified splice segments in all three species. From these nine splicing events, eight showed consistent differences in isoform concentrations among three primates coinciding with the differences observed in RNA-seq data in all three biological replicates (Figure 6B, C; Supplementary Figure S5; Supplementary Table S5).

Among these eight splicing events, five were predicted to affect the protein sequence of the gene (Supplementary Figure S5). To test whether these splicing changes led to the expression of different protein isoforms in humans compared to chimpanzees and macaques, we performed Western blot experiments using cerebellum samples from three humans, three chimpanzees and three macaques (Supplementary Table S6). From the four antibodies tested, only the polyclonal antibody against CEP63 protein produced specific bands. In agreement with our predictions based on the RNA-seq results, the Western blot showed the presence of one CEP63 protein isoform in the chimpanzee and macaque brains and two protein isoforms in the human brain (Figure 6D-F).

To assess the potential functional significance of the identified species-specific splicing changes, we tested the enrichment of splicing events from D1, D2, and I2 splicing patterns among the Gene Ontology (GO) functional categories. Among all lineages and patterns, the human-specific I2 and D2 patterns showed significant functional enrichment (FDR<0.05; Figure 6G, H; Supplementary Table S7, S8). Interestingly, although the enriched GO terms for I2 and D2 patterns did not overlap, in both cases they were mainly related to neuronal functionality. The D2 pattern was enriched in seven cellular component GO terms predominantly associated with synaptic location, while the I2 pattern was enriched in 23 biological process GO terms including axon guidance and related functions. Functions enriched in the I2 pattern were reported to be affected by alternative splicing of cell adhesion proteins (28–30). These proteins include products of two cell adhesion genes from the L1 family, CHL1 and NRCAM, and two semaphorin genes, SEMA4D and SEMA6C, which we identified to be alternatively spliced in our study. More generally, among 33 genes containing I2 or D2 splicing changes on the human lineage and falling into the enriched GO terms, 29 showed significant splicing changes in at least one brain region, and 19 included splicing changes that altered protein sequences.

## DISCUSSION

Splicing evolution includes a variety of processes, including birth and death of novel isoforms, as well as increases and decreases in the frequency of existing ones. We investigated each of these processes separately by classifying differences in exon inclusion levels among four species into six evolutionary patterns. This classification resulted in an unexpected observation: different splicing patterns substantially varied in their frequencies and potential functional implications.

Among the six splicing patterns, one representing the birth of novel isoforms by constitutive-to-alternative exon transition contained 60-65% of all detected splicing differences. Despite its prevalence, our results suggest that this pattern represents neither the primary driving force of long-term isoform repertoire evolution nor the source of novel gene functionality. Isoforms created within this pattern often result in reading frame disruption, are less conserved at the DNA sequence level, do not cluster in any functional categories, and are lost more rapidly during the course of evolution than isoform changes in the other splicing patterns. Our analysis also suggests that constitutive-to-alternative splicing changes might be primarily driven by mutations within proximal splice sites and splicing factor binding motifs, thereby decreasing their efficiency. Overall, these observations strongly imply the neutral or deleterious nature of splicing events leading to constitutive-to-alternative exon transition. Nonetheless, we cannot exclude that perhaps a few of the novel alternative isoforms contained in this pattern may gain new functionality during evolution.

The majority of the remaining splicing events (22-27% of the total) does not affect isoform numbers but instead changes their relative frequencies. These alternative isoforms tend to preserve protein reading frames, are highly conserved at the DNA sequence level, but tend to show high Ka/Ks ratios indicating an acceleration of protein sequence evolution, and tend to be preserved by evolution. We further identified mutations in trans- regulator binding sites as a potential regulatory mechanism driving this type of splicing change. These observations suggest that changes in inclusion frequencies of existing isoforms represent functional evolutionary transitions.

Changes in the inclusion frequency of existing isoforms might be particularly relevant for evolution of the human brain. One of the two patterns constituting this group of events, the increase in isoform inclusion levels, showed a two-fold acceleration on the human evolutionary lineage when compared to the chimpanzee lineage. This acceleration particularly affected the isoform repertoire in the brain. Furthermore, despite the relatively small number of observed splicing events, both patterns clustered significantly in biological processes relevant to brain function.

Due to the limited numbers of detected events, we were unable to provide a detailed characterization of the remaining three evolutionary splicing patterns. However, this omission does not mean that the splicing processes described by these patterns do not play a role in evolution. For instance, novel exon birth is one of the essential mechanisms leading to the emergence of new gene functions (31–33). As larger data sets representing a wider selection of species and evolutionary distances appear, categorizing splicing events into discrete evolutionary patterns might lead to novel insights into the evolutionary mechanisms and functional consequences of splicing changes.

## MATERIALS AND METHODS

### Samples

The RNA-seq data used in this study was taken from (19) (Ensembl ID: GSE49379, Supplementary Table S1). Selected human, chimpanzee, and rhesus macaque samples used in the RNA-seq experiment were also utilized in the PCR validation experiments, as labeled in Supplementary Table S1.

For Western blot, human samples were obtained from the National Institute of Child Health and Human Development (NICHD) Brain and Tissue Bank for Developmental Disorders at the University of Maryland. Written consent for the use of human tissues for research was obtained from all donors or their next of kin. All subjects were defined as healthy controls by forensic pathologists at the corresponding tissue bank. All subjects suffered sudden death with no prolonged agonal state. Chimpanzee samples were obtained from the Biomedical Primate Research Centre, the Netherlands. Rhesus macaque samples were obtained from the Suzhou Experimental Animal Center, China. All nonhuman primates used in this study suffered sudden deaths for reasons other than their participation in this study and without any relation to the tissue used. The cerebellum samples were dissected from the lateral part of the cerebellar hemispheres.

### Reads mapping

1.87 million 1×100nt single-end RNA-seq reads were mapped to human, chimpanzee, rhesus macaque, and mouse reference genomes (genome assemble version: hg19, panTro3, rheMac2, and mm9) using the RNA-seq read mapping software Tophat (v2.0.3, http://tophat.cbcb.umd.edu/) (34). In the first round of mapping, we detected de-novo junctions in each sample. A total of 1.34 billion reads, including 305 million junction reads, were successfully mapped to reference genomes (mapped proportion: 72%, Supplementary Table S2). We combined all detected junctions after we transferred their coordinates to other species using liftOver (https://genome.ucsc.edu/cgi-bin/hgLiftOver). The union of all junctions was inputted in the second round of mapping in order to equalize junction detection sensitivity in all species. Considering that human genome annotations are more complete than those for chimpanzees and macaques, here we did not use public gene structure annotations in order to avoid analysis bias.

### PSI calculation

To minimize interspecific artifacts caused by length or mapping efficiency changes inside exons, we only used junction reads for the PSI value calculation. The PSI value is equal to the proportion of inclusion reads among the total reads. For example, let [*L*, *R*] denote the exclusion junction, *i.e.* the junction supporting the exon-exclusive isoform, where *L* and *R* represent the positions of its left (5’-side) and right (3’-side) splice sites respectively. Also, let [*L, M*_*1*_] and [*M*_*2*_, *R*] denote its left and right inclusion junctions, *i.e.* the junction supporting the exon-inclusive isoform, where *M*_*1*_ and *M*_*2*_ are two reference positions between *L* and *R*. In the event of a single cassette exon, *M*_*1*_ and *M*_*2*_ are the two ends of this exon. For each species, the PSI value of this gene segment (one or several adjacent cassette exons) in tissue *t* was calculated as follows:

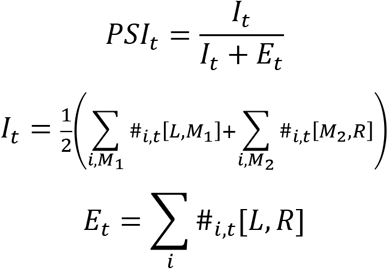

where #_*i,t*_[*L, M*] denotes the observed read number of this junction in individual *i* and
tissue *t*, and *I*_*t*_ and *E*_*t*_ denote the inclusion and exclusion junction read number respectively. Lastly, only splicing events that satisfied the criteria below were used for further analysis:

1. Must have at least one inclusion or exclusion junction read in every 90 primate individuals for three-species comparison analysis, or one read in every 120 individuals for four-species comparison analysis;
2. In at least one species-tissue combination, there must not be less than 4 individuals with a left inclusion read, and also no fewer than 4 individuals with a right inclusion read; in at least in one species-tissue combination, there must not be less than 4 individuals that have an exclusion read;
3. Both **Σ**_*i*_#_*i,t*_[*L*,*M*_1_ and **Σ**_i_#_*i,t*_[*M*_2_, *R*] should be larger than the number of non junction reads covering the left or right splice site, *i.e.*, the retention ratio of either the left or right intron should be less than 0.5; **Σ**_i_#_*i,t*_[*L*, *M*_*1*_] should not be less than 15% of **Σ**_i_ #_*i,t*_[*M*_*2*_, *R*], and *vice versa*. This criterion aims to exclude events with obvious intron retention, alternative donor sites or alternative acceptor sites;
4. Both for inclusion and exclusion reads, the multiple-mapped read number should be less than 15%. This criterion can alleviate the bias that arises when a splicing region has a unique nucleotide sequence in the genome of one species but is duplicated in the genome of another species.

### Detecting lineage-specific spliced gene segments

We first required that the splice sites of analyzed cassette exon sets should be conserved among all compared species; and that the genes should be expressed in all individuals of these species. In the primate comparison assay, rhesus macaque was regarded as the outgroup in order to detect splicing changes that occurred on the human and chimpanzee lineages. In the four-species comparison assay, mouse was regarded as the outgroup in order to detect splicing changes that occurred on the ape and rhesus macaque lineages. All comparisons were done separately in each tissue.

In the primate comparison assay, a quasi-binomial generalized linear model was applied based on inclusion and exclusion read numbers, using the glm function in R. Chi-squared test p-values were calculated using the species term of this model. For multiple-test correction, 1,000 permutations were performed by randomly shuffling the species labels. We used 0.05 as the false discovery rate cutoff. We further required that the differences between the maximum and minimum PSI values among the three primate species be larger than 0.2. The cassette exon sets that met these criteria were considered as differentially spliced in primates. Next, we classified each differentially spliced exon set as human lineage-specific, chimpanzee lineage-specific, or other, in each of the five tissues. For example, we regarded min(|*H* − *C*|, |*H* − *R*|) > 2|*C* − *R*| as human lineage-specific, and regarded min(|*H* − *C*|,|*C* − *R*|) > 2|*H* − *R*| as chimpanzee lineage-specific, where H, C, and R denote the PSI values in human, chimpanzee, and rhesus macaque, respectively.

In the four-species comparison assay, we also used the quasi-binomnial GLM test to detect divergent cassette exon sets, followed by 1,000 permutations for multiple-test correction with a 0.2 PSI change cutoff. To further detect splicing changes specific to the rhesus macaque and ape lineages, we compared the PSI values of mouse (*M*), macaque (R), and ape (*i.e.* the mean value of human and chimpanzee) (*A*). We required that the inclusion level on the mouse lineage should be consistent with the inclusion level either on the macaque or ape lineages (*i.e.* |*M* − *A*| < 0.2 or |*M* − *R*| < 0.2). Under this condition, cassette exon sets that satisfied min(|*M* − *A*|, |*R* − *A*|) > 2|*M* − *R*| were regarded as ape lineage-specific, while the cassette exon sets that satisfied min(|*M* − *R*|, |*R* − *A*|) > 2|*M* − *A*| were regarded as rhesus macaque lineage-specific. In the birth-and-death analysis of splicing patterns (Figure 3 B-D), equivalent criteriamin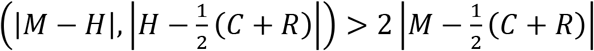 and 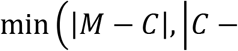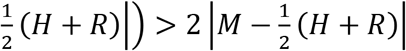 were applied to human and chimpanzee lineage-specific cassette exon sets in order to make the numbers of changed cassette exon sets comparable among the four lineages.

For the convenience of comprehensively analyzing splicing changes, we chose a representative tissue from the five tissues for each changed cassette exon set. The representative tissue chosen had the smallest p-value of interspecific PSI change, among the tissues whose splice change lineage assignment was the most common. For example, if a cassette exon set has *p*-values {0.01, 0.005, 0.03, 0.05, 0.05} in {muscle, kidney, prefrontal-cortex, visual-cortex and cerebellum} tissues, specifically spliced in {human, macaque, human, human, unassigned} lineages respectively, then the representative tissue is muscle, since its most common splice change lineage assignment is human and muscle has the lowest *p*-value among them (0.01).

The remaining conserved cassette exon sets with PSI changes below 0.2 among three primates in all tissues were used as a control group on further analysis.

### Defining the six splicing change patterns

We compared PSI values of lineage-specific cassette exon sets in the changed lineage against the other two (primate assay) or three (four-species assay) unchanged lineages. For each exon set, if its PSI value on the changed lineage was below 0.03 or above 0.97, it was assigned to the D3 or I3 pattern respectively; if its PSI value on any of the unchanged lineages was below 0.03 or above 0.97, it was assigned to the I1 or D2 pattern respectively; otherwise, it was assigned to the I2 or D2 pattern depending on the direction of change.

### Birth and death rate calculation for the D1 and I2 + D2 patterns

Suppose that in a unit of evolutionary time, *e.g.* a million years, the probability of a cassette exon set whose splicing changes in a specific pattern (pattern birth) is *p*_*1*_; and for a cassette exon set whose splicing formerly changed, the probability of a reverse change (pattern death) is *p*_*2*_. For the human, chimpanzee, and rhesus macaque lineages, after a time period *t*, the expected cassette exon set number changed on lineage *s* (*m*_*s*_) in a given pattern can be described by the differential equation below:

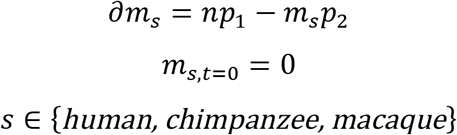

Where *n* is the number of total alternatively spliced cassette exon sets. Solving this equation, we get the proportion *q* of cassette exon sets with detectable splicing changes as a function of *t*, as below:

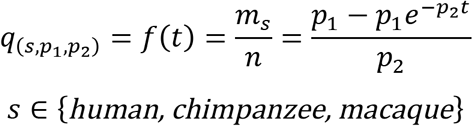

For simplicity we did not consider the case of parallel evolution, as its probability is extremely low.

Next we built a model for the ape lineage. Let *t*_*1*_ denote the time range of the ape lineage (the green line in Figure 3A), and *t*_*2*_ denote the time range since the human and chimpanzee lineages separated. A pattern specific to the ape lineage can both be born and die during *t*_*1*_, but can only die during *t*_*2*_, since pattern death occurring in either the human or chimpanzee lineages would make the ape-specific change undetectable. The expected number of cassette exon sets specifically spliced in the ape lineage *m*_*ape*_ can be described according to the differential equation below:

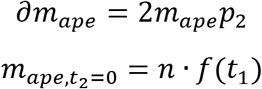

Solving this equation, we also get the proportion *q* of cassette exon sets with splicing changes detectable in the ape lineage as:

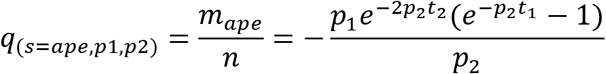

For all four lineages, it is reasonable to assume that the actual observed number of cassette exon set changes *M*_*s*_ in lineage *s* follows a binominal distribution:

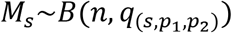

Therefore, the likelihood for given*p*_*1*_ and*p*_*2*_ is:

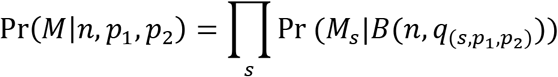

To find *p*_*1*_ and *p*_*2*_ with the highest likelihood, we calculated Pr(*M*|*n*, *p*_1_, *p*_2_) in a 1000 × 1000 Cartesian space, in the range of *p*_*1*_=[0, 0.003], *p*_*2*_=[0, 0.1] for the I2+D2 patterns, and *p*_1_=[0, 0.015], *p*_*2*_=[0, 0.25] for the D2 pattern. As the likelihood for *p*_*1*_ and *p*_*2*_ in the boundaries of these spaces was extremely small, we ignored the likelihoods outside of this area. In Figure 3D we also show the high-likelihood sub-areas for the confidence interval. The likelihoods of these sub-areas are 95% and 99% of the sum of the total likelihood respectively.

### Ka/Ks analysis

For this analysis, we used coding regions of gene segments specifically spliced on human and chimpanzee lineages. The sequences of these regions were compared to a primate consensus genome in order to find out the nucleotide mutations on either the human or chimpanzee lineages. The numbers of synonymous and non-synonymous mutations were calculated using the program yn00 in PLAM (v4.7a, http://abacus.gene.ucl.ac.uk/software/paml.html) (35).

### Splice site analysis

We used Splice Site Tools (http://ibis.tau.ac.il/ssat/SpliceSiteFrame.htm) to assess the strength of splice sites. The 5’ splice site strength was calculated based on nine nucleotides from position -3 to +6, while the 3’ splice site strength was calculated based on 15 nucleotides from position −14 to +1. We only used splice sites flanking cassette exon sets with consistent strengths between human and chimpanzee lineages, and compared them to the rhesus macaque lineage. We used a total of 128 D1 exon sets specifically spliced on either the rhesus macaque or ape lineage, which were detected in the four-species comparison. To get sufficient I2 + D2 exon sets for statistical analysis, we used 178 cassette exon sets, which were alternatively spliced in all three primates (0.03<PSI<0.97), with similar inclusion levels in the human and chimpanzee, but different inclusion levels from the macaque (assessed using similar criteria as above). These exon sets had I2 and D2 splicing changes on either the rhesus macaque or ape lineages. The other 1,026 cassette exon sets with PSI changes below 0.2 among three primates were used as a control group.

### RNA binding motif analysis

Sequences of 329 RNA binding motifs from 52 splicing factors in humans were downloaded from SpliceAid database (http://www.introni.it/splicing.html) (36). To check for the existence of these motifs in the three primate species, we screened six regions of each cassette exon, including 200nt from the proximal and distal ends of adjacent introns (with an intron length >300nt) and 39nt from the ends of the cassette exon (with an exon length >60nt). We detected motif changes (turn-on and turn-off) on both the human and chimpanzee lineages using rhesus macaque as an outgroup.

### GO enrichment analysis

We tested GO enrichment for each combination of the three dominant patterns (D1, I2, and D2) and three primate lineages. Genes with specific splicing changes in the tested dominant patterns and primate lineages in any of five tissues were used. For the I2 and D2 patterns, we excluded marginal cases by requiring that the minority isoform in any primate species should be greater than ten percent. For each test, a background gene set was randomly chosen from the cassette-exon-containing genes, to make sure that the background gene set had a similar expression distribution as the test gene set. GO enrichments were tested using the R package topGO (37) with “classic” algorithm and “Fisher exact test”, followed by the Benjamini-Hochberg multiple test correction with significant cutoff FDR=0.05. To make the results more concise, if two GO terms from father and child nodes were both significant with the exact same matched genes, only the child term was reported. The enriched ontologies and their significances are shown in Supplementary Table S7, S8.

### PCR confirmation

PCR primers were designed at the two neighboring constitutive exons of spliced gene segments in order to detect longer (exon-included) and shorter (exon-excluded) products (Supplementary Table S5). First-strand cDNA was synthesized using SuperScript II reverse transcriptase (Invitrogen). After double strand cDNA synthesis, PCR reactions were performed at 94°C for 2min followed by 30 cycles of 95°C for 20s, 47~58°C (depending on the primers) for 20s, and 72°C for 30s, and complete elongation at 72°C for 5 min with rTaq DNA polymerase (TAKARA). Each PCR product was verified in a 2% (w/v) agarose gel, with size estimation based on the DL 2000 DNA marker (Takara), or verified by DNA 1000 chip (Agilent) as the protocol suggested. The PCR PSI values were calculated as the mean proportion of the longer product strength between the two expected products. To validate primer efficiency in all species, we first tested twelve cassette exon sets in three PCRs (3 species × 1 tissue × 1 individuals). Nine produced discernable PCR bands with sizes corresponding to two predicted alternative transcript isoforms. Out of the nine cases, one showed inconsistent band intensities in the rhesus macaque for unknown reasons. The other eight cases were further expanded for a total of nine PCRs (3 species × 1 tissue × 3 individuals). All repeat experiments showed band intensities consistent with our RNA-seq results (Supplementary Figure S5).

### Western blot

Western blot samples are in Supplementary Table S6. Among the seven tested antibodies, Anti-CEP63 polyclonal rabbit antibody was purchased from Merck Millipore (Catalogue Number: 06-1292). Electrophoresis and western blot procedures were carried out as described in the NuPAGE Novex system (Invitrogen). Briefly, the protein was denatured at 70°C for 10 min with reduced loading buffer. The electrophoresis cassette was set and run at 200V for 1-1.5 h according to the size of the target protein. The blotting sandwich was assembled and the protein was transferred from the gel onto the polyvinylidene difluoride (PVDF) membrane at 100V for 1-2 h. After unspecific binding was blocked for 2h at room temperature, the membrane was incubated with the primary antibody at 4°C overnight. The next day, the membrane was incubated in horseradish peroxidase (HRP)-linked secondary antibody for 1 h after a brief wash in PBST (PBS with 0.5% TWEEN 20). The chemiluminescent signal was generated by adding the substrate to the membrane and exposing it with photographic film in the dark room.

